# InteBac - An integrated bacterial and baculovirus expression vector suite

**DOI:** 10.1101/2020.07.09.194696

**Authors:** Veronika Altmannova, Andreas Blaha, Susanne Astrinidis, Heidi Reichle, John R. Weir

## Abstract

The successful production of recombinant protein for biochemical, biophysical and structural biological studies critically depends on the correct expression organism. Currently the most commonly used expression organisms for structural studies are *E. coli* (ca. 70% of all PDB structures) and the baculovirus/ insect cell expression system (ca. 5% of all PDB structures). While insect cell expression is frequently successful for large eukaryotic proteins, it is relatively expensive and time consuming compared to *E. coli* expression. Frequently the decision to carry out a baculovirus project means restarting cloning from scratch. Here we describe an integrated system that allows the simultaneous cloning into *E. coli* and baculovirus expression vectors using the same PCR products. The system offers a flexible array of N- and C-terminal affinity, solublisation and utility tags, and the speed allows expression screening to be completed in *E. coli*, before carrying out time and cost intensive experiments in baculovirus. Importantly, we describe a means of rapidly generating polycistronic bacterial constructs based on the hugely successful biGBac system, making InteBac of particular interest for researchers working on recombinant protein complexes.

## Introduction

Obtaining recombinant protein of interest can be a challenging multi-parametric problem. The parameters to consider include codon usage, vector, fusion tag, expression organism, expression conditions and purification strategy^1^. Previous work has described the use of universal vectors compatible with *E. coli*, baculovirus and mammalian expression systems for example the pOPIN system^2^. However for insect cell expression the excellent MultiBac system^3^ has set the standard. A recent baculovirus expression method has combined MultiBac with Gibson assembly^4^ to yield the biGBac system^5,6^. Using biGBac one can assemble a five open reading frame polycistronic vector in a single step, and combine five of these in a second step to assemble up to 25 ORFs in a single vector. Our overwhelming satisfaction with biGBac lead us to further develop the system while creating a parallel, compatible, system for *E. coli*.

We have previously used the pST44 polycistronic expression system for bacterial expression^7^. Not only does pST44 provide a rapid means of generating multicistronic constructs, but the pST44 familiy pTRC50 vectors are excellent expression vectors in their own right. However, the pST44 method uses restriction enzyme based approaches which has been superseded by ligation independent approaches, particularly Gibson assembly, in the last 10 years^4^.

Until recently in our lab, one would make a decision to pursue either a *E. coli* approach or an insect cell approach to obtaining a recombinant protein of interest. Working in parallel was of course always possible, but would require creating different PCR products, preparing different vectors, and being limited by the fusion proteins available for each system. To address this limitation we set out to create a unified and integrated bacterial/insect cell expression system. Our goal was to be able to take a single PCR product and clone this into numerous *E. coli* and insect cell expression vectors. One would process both sets of vectors in parallel, and have the result for *E. coli* expression before one even transfected insect cells. The ultimate result being that one has more time and resources to explore the parametric space of recombinant protein expression (summarised in Figure 1).

**Fig. 1.**
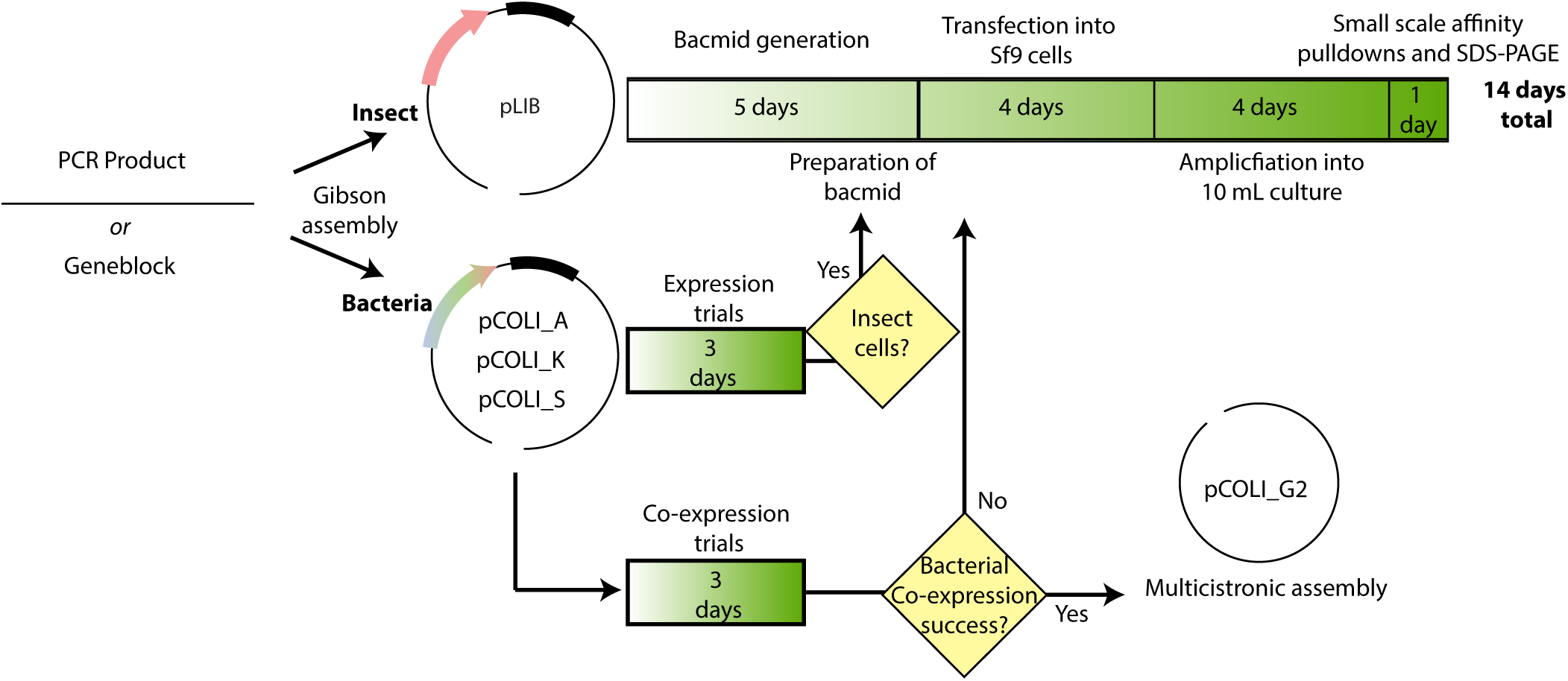
Workflow within the InteBac and biGBac systems. Using a single PCR product one clones simultaneously into pLIB (for insect cells) and pCOLI (for bacteria). Depending upon the results of the *E. coli* expression trial, one can decide whether or not to proceed with insect cell work.

**Fig. 2.**
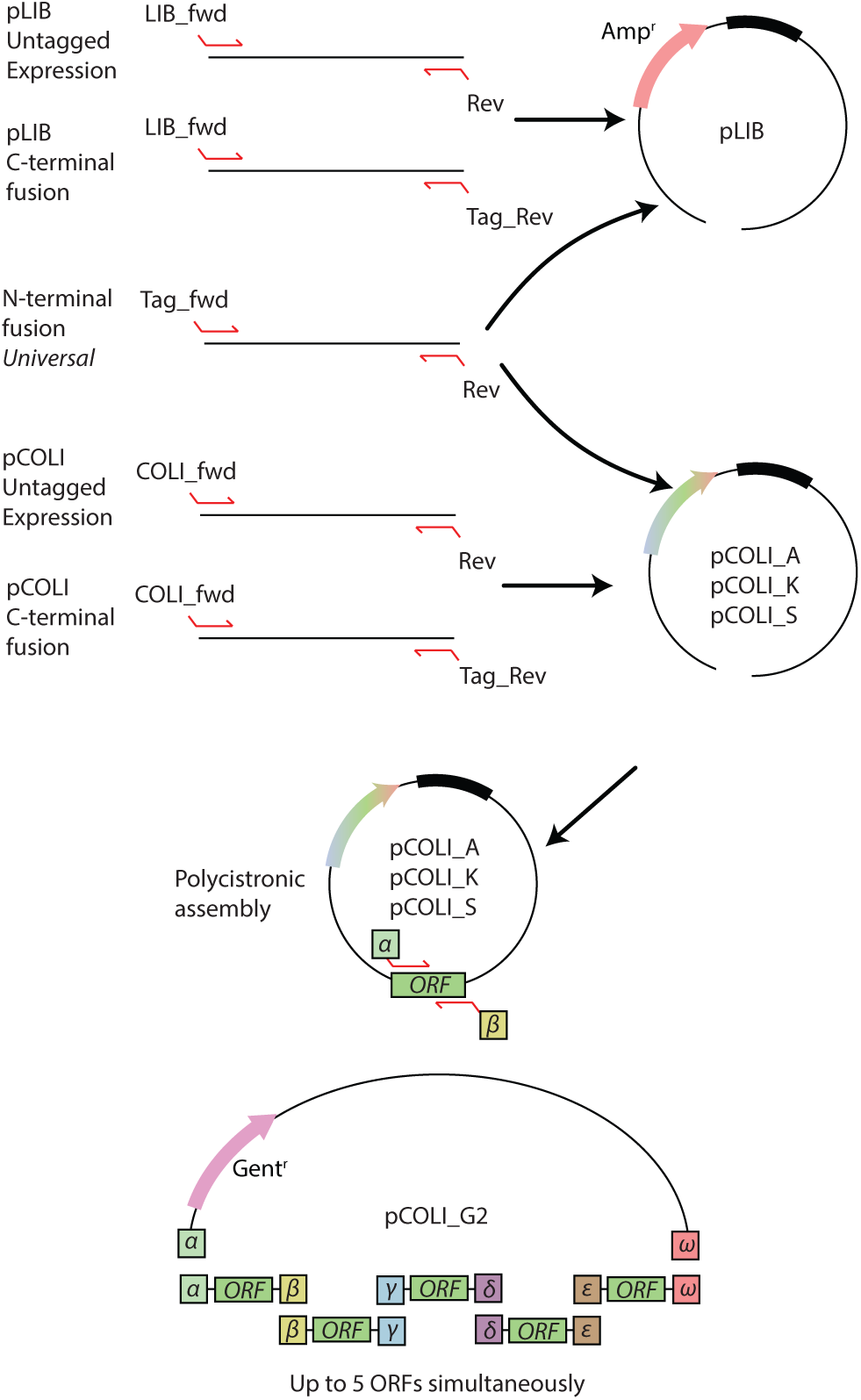
Cloning in the InteBac system. Primers, or overhangs on geneblocks, are chosen to match the vector. The N-terminal fusion vectors are truely universal, allowing for cloning into either the pLIB or the pCOLI backbones. From the pCOLI backbones one can generate a polycistronic construct with up to five insertions. Fewer insertions can be used, but the alpha and omega overhangs must be present.

When pursuing the expression of multi-subunit complexes, the creation of a polycistronic construct, for either insect cell or bacterial expression, is a useful means of both ensuring appropriate complex stoichiometry but also reducing the complexity of a biochemical reconstitution. However, the establishment of the appropriate conditions by screening subunits and fusion tags is easier when combining multiple monocistronic constructs in co-expression experiments. In insect cells, this can be achieved by co-infecting with multiple different viruses, albeit with some limitations (described in 8). We seldom co-infect with more than two viruses, and it is important to screen different ratios of virus. In our lab this is usually done volumetrically combining different ratios of virus A with with virus B. In *E. coli*, the situation is complicated by both antibiotic and origin of replication usage and subsequent plasmid incompatibility. To this end, we modified our *E. coli* vector set with origins of replication and antibiotic resistances compatible with co-expression.

Here we describe the implementation of a parallel system for the rapid screening of *E. coli* co-expression vectors that are compatible with simultaneous cloning into insect cell expression vectors. Furthermore we describe a Gibson assembly based approach for the rapid generation of polycistronic *E. coli* expression constructs. This system has greatly improved the workflow in our lab, and we hope other labs will benefit from our efforts.

## Results

### Generation of N- and C-terminal fusion vectors

We started with creating an insect cell expression vector based on the pLIB vector^5,6^ with a variety of N-terminal fusion proteins. We divided the tags into different categories; affinity, solublisation, and utility (see Table 1). In order to have universal overhangs for both the insect cells and bacterial expression systems, these were designed to correspond to a rhinovirus-3C site between the fusion protein and the protein of interest (see Table 2 for all primer overhangs). We chose 3C cleavage site over the more frequently used TEV site, due to the 3C protease’s higher catalytic activity at lower temperatures^9^. Next we created a more limited set of C-terminal fusion proteins, placing a serine-glycine linker between the protein of interest and the C-terminal fusion protein. This linker ensured that the C-terminal overhang would be universal for all C-terminal fusions. We transferred these fusion protein ORFs from the pLIB backbone into the pTRC50 backbone (Ampr / pBR232 origin^7^), for bacterial expression (from now on referred to as pCOLI_A). Finally, in order to give us the greatest flexibility in *E. coli*, we transferred all the expression cassettes into two additional backbones pCOLI-S (Strepr/RSF1030 origin^10^) and pCOLI_K (Kanr / CloDF13 origin^11^). This combination of resistances and origins of replication gives the user the ability to co-express three proteins simultaneously (see Supplementary Table 1 for exhaustive list of all expression vectors).

**Table 1.** Summary of fusion proteins used in the InteBac system.

| Fusion name | Description | Type | Mw N-term fusion |
| --- | --- | --- | --- |
| 6xHis | IMAC purification | Affinity | 3 kDa |
| 12xHis | IMAC from insect cells |  | 3.8 kDa |
| STREP | Twin Strep-II tag |  | 4.9 kDa |
| CBP | Calmodulin binding peptide |  | 4.3 kDa |
| GST | Glutathione-S-transferase | Solubilisation /Affinity | 26.7 kDa |
| MBP | 6x His plus Maltose binding protein |  | 42.7 kD |
| SUMO | 6 x His plus SUMO | Solubilisation | 13.5 kDa |
| Trx | 6 x His plus Thioredoxin |  | 14 kDa |
| SNAP | 6 x His plus SNAP tag | Utility | 20 kDa |
| HA | 6 x His plus 3 x HA | Identification | 6.3 kDa |
| Myc | 6 x His plus 6 x Myc |  | 11 kDa |

**Table 2.**
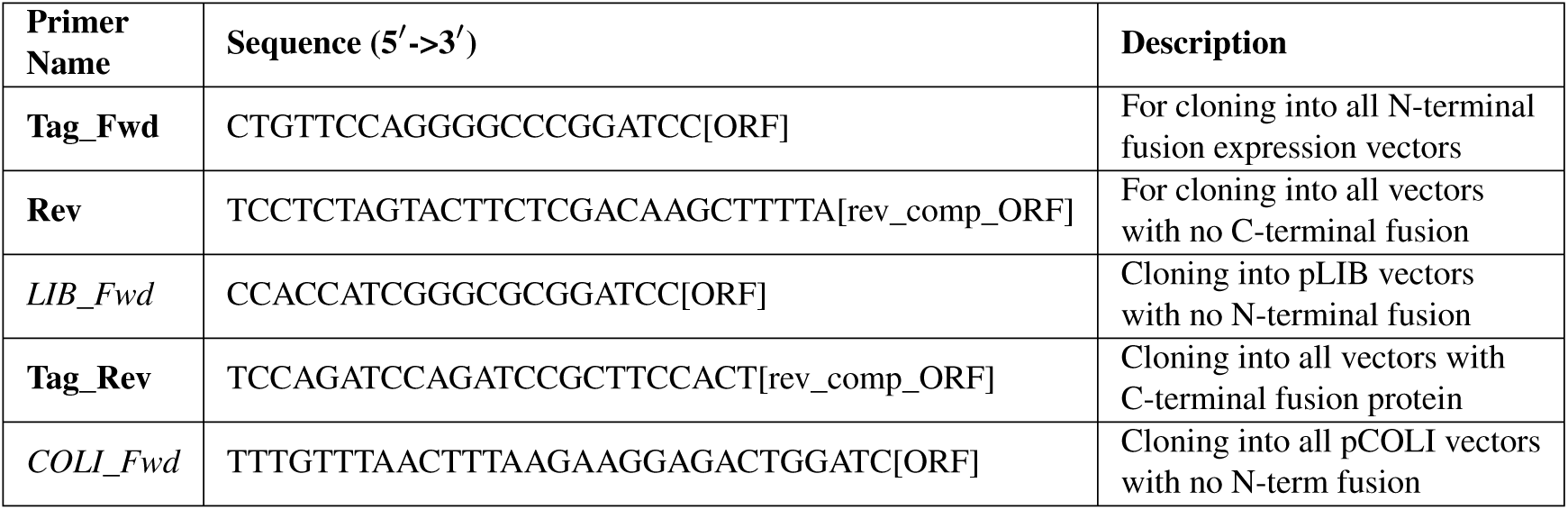
Primers used to clone the gene of interest into the pLIB and pCOLI vectors as untagged or C- or N-terminal fusion constructs.

### Untagged vectors

Many proteins are not amenable to N- or C-terminal tagging, but can be purified through the affinity tag on a binding partner. In order to facilitate co-expression of several proteins we required untagged vectors. Despite our efforts we were unable to generate a generic N-terminal overhang that would work for both untagged *E. coli* and insect cell vectors. As such, there is a generic overhang for untagged insect cell and untagged *E. coli* vectors, rather a specific forward primer for each (pLIB_fwd and COLI_fwd respectively, see Table 2). To facilitate co-expression in *E. coli* we also created untagged pCOLI_S and pCOLI_K vectors. These vectors contain compatible origins of replication and resistances to facilitate co-transformation into bacteria.

### Multicistronic vectors

Our pLIB derived vectors remain fully compatible with the pBIG multicistronic vectors from the biGBbac system^5,6^. To create a multicistronic bacterial vector we took the pST44 vector backbone and added a gentamycin cassette (from now on referred to pCOLI_G2). We designed a set of PCR primers for amplifying the entire ORF from the pCOLI family of vectors (including the RBS, but not the promotor or terminator). The Gibson overhangs described in the biGBac system were thoroughly tested, both *in silico* and *in vitro*, to give the greatest assembly efficiency. As such we use the same principle, and indeed the same overhang sequences as in biGBac, to create at multicistronic pST44 vector, in addition to the use of the SwaI enzyme.

### Proof of concept - RPA complex

Our interest in homologous recombination has led us to look at several protein complexes involved, one of which is RPA (Replication Protein A, reviewed in 12&13), a heterotrimer consisting of Rfa1, Rfa2 and Rfa3 in budding yeast^14^. Expression and purification of yeast RPA in E. coli has been previously described^15^.

We cloned Rfa1, Rfa2 and Rfa3 into Strep-pCOLI-A, His-pCOLI-S and His-pCOLI-K respectively. We initially demonstrated that the complex could be expressed and partially purified through a co-expression of all three proteins in *E. coli* C41(DE3), and purification via the twin Strep-II tag, followed by confirmation of protein identity via western blotting (Figue 3, lanes 1-4). We amplified the expression cassettes for each of the three RPA subunits, and Gibson assembled into the linearised pCOLI-2G backbone. Gentamycin resistant transformants were confirmed by sequencing, and subsequently transformed into the BL21 cells.

**Fig. 3.**
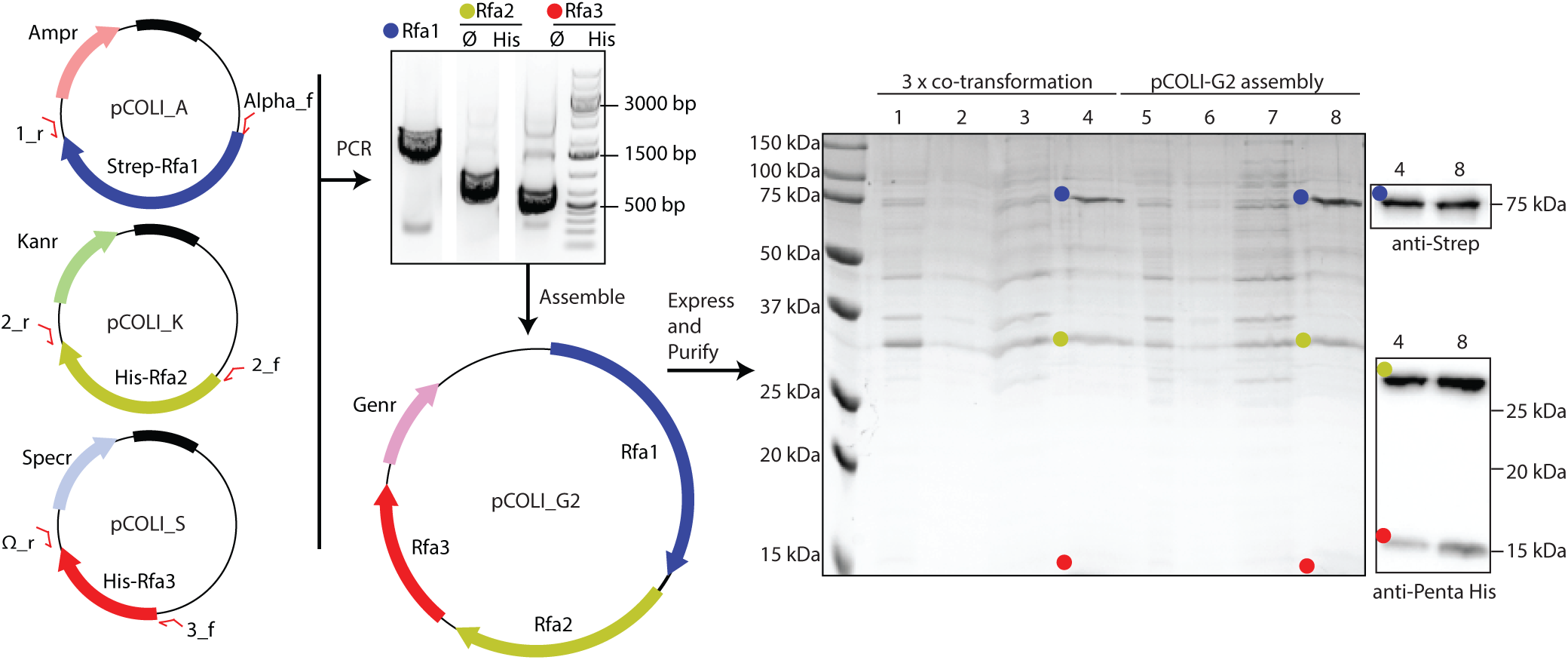
Cloning and expression of the trimeric RPA complex from yeast. Each RPA subunit was cloned into a different pCOLI backbone, which were then used for co-expression. Additionally we generated a multicistronic assembly of RPA into the pCOLI_G2 backbone. We compared the expression of the co-tranformation versus pCOLI_G2. SDS-PAGE was run of crude lysate (lanes 1 and 5), clear lysate (lanes 2 and 6), flow through from the resin (lanes 3 and 7), elution from the beads (4 and 8)

Our Gibson assembly of polycistronic RPA was just as successful, if not more so, than the co-expression of all three RPA subunits. Furthermore there is the advantage of carrying out transformations with a single plasmid, using a single selection antibiotic, and the possibility of carrying out further co-expressions with pCOLI-K and pCOLI-S (Figure 3).

## Discussion

Our system is also well suited to the use of “oven-ready” synthetic dsDNA (Geneblocks), allowing the incorporation of the Tag_fwd and Rev overhangs into the synthetic DNA, prior to assembly into the pLIB or pCOLI vectors. Currently such synthetic dsDNA fragments are available up to ca. 3000 bp in length, though this usually requires screening multiple colonies to find a transformant with the correct sequence. Sequence verified dsDNA fragments are also available, but at a higher price. Additionally we would recommend users to explore the use of codon optimisation. In our experience constructs optimised for expression in *E. coli* K12 also work well in either *S. frugiperda* or *T. ni* (Sf9 and Hi5 cells).

The implementation of our InteBac system has greatly streamlined the work processes in our laboratory. It has allowed us to explore additional experimental space in terms of finding suitable expression conditions for our protein of interest. Previous systems, including the “pCoofy”^16^ have also made use of single PCR products to be integrated into a range of different expression vectors; including those for insect cell expression. InteBac differs from pCoofy in several key areas. Firstly, InteBac has been designed to work as an “add on” to the biGBac system, allowing the generated insect cell expression vectors to be turned into multicistronic vectors through a single Gibson assembly step. Secondly, InteBac makes use of three different bacterial expression backbones facilitating rapid co-expression of up to three proteins in *E. coli* before moving to a polycistronic construct. Finally, we offer the possibility of producing a multicistronic *E. coli* expression vector (based on the principles outlined in biGBbac ^5,6^) in pCOLIG2. As such we consider InteBac of particular interest to those researchers working on protein complexes.

## Materials and Methods

### Vector construction

All cloning and plasmid manipulation steps were first carried out *in silico* using the SnapGene software (GSL BioTech LLC). The pTRC50 and pLIB vectors were gifts from Song Tan (Penn State) and Jan Michael Peters (IMP Vienna), respectively. PCR amplifications were carried out using 2 x Q5 Master Mix (NEB), with cycling times and temperatures according to the manufacturer’s instructions. The Kanamycin and Streptamycin resistance/origin of replication modules were synthetic dsDNA constructs (IDT). The gentamycin cassette was amplified from the pBIG2 vectors. Since the gentamycin cassette also contains one restriction site for BglII, we introduced a silent mutation into its sequence. All affinity tags insertions and plasmid manipulation was carried out using a combination of synthetic dsDNA (IDT) and Gibson assembly. Successful assemblies were verified by Sanger sequencing.

### Gibson Master Mix

For all Gibson assemblies we used our own master mix. Briefly, a 5 x isothermal reaction buffer was prepared (25% PEG-8000, 500 mM Tris-HCl pH 7.5, 50 mM MgCl2, 50 mM DTT, 1 mM each of the 4 dNTPs, and 5 mM NAD) and pre-aliquioted. To prepare Gibson master mix we combined 320µl 5X ISO buffer with 0.64 µl of 10 U/µl T5 exonuclease, 20 µl of 2 U/µl Phusion polymerase and 160 µl of 40 U/µl Taq ligase (all enzymes from NEB). ddH2O was then added to a final volume of 1.2 ml.

### Linear Vector Preparation

All vectors (with the exception of pCOLI_G2) are designed to be linearized with the same combination of restriction enzymes (BamHI and HindIII). Proper vector linearization and subsequent purification is critical to the success of downstream cloning. Briefly, we took 1 µg of plasmid (typically from a midi-prep (Qiagen)) and digested in a 20 µL reaction with 20 units of BamHI-HF (NEB) and 20 units of HindIII-HF (NEB) in CutSmart buffer (NEB) for 3 hrs. Each reaction was then gel purified using the Wizard SV kit (Promega) according to the manufacturer’s instructions, with the exception that 30 µL of ddH2O was used for elution from the column. The final linearized plasmid product had a typical concentration of 60-100 ng/µL. To linearize pCOLI_G2 vector the restriction enzymes BglII and XhoI were used and the plasmid was further processed as described before.

### Insert preparation

All inserts were amplified using Q5 polymerase (NEB). PCR reactions were gel purified using Wizard SV gel and PCR cleanup. In case of amplification from a plasmid source, we paid attention to the size of the insert versus the template. In case of any potential overlap we treated our PCR reactions with 1 µL DpnI (NEB) to eliminate the donor plasmid. DpnI was then heat inactivated (15 minutes 65°C) prior to gel purification.

### Gibson cloning and verification

Routinely we mixed 4.5 µL of purified insert with 0.5 µL of vector and added this 5 µL to one 15 µL aliquot of Gibson master mix. Our Gibson reactions were then incubated for 1 hour at 50°C, and then transformed directly into chemically competent XL1-Blue. From a large number of colonies we would typically grow two, and prep one for sequencing, keeping the other as a backup. Typically, with a well-prepared vector (see above) our cloning success rate is >95%, so we considered it wasteful to “pre-screen” with analytical digests. All agarose gels shown are 0.8% agarose, stained with GelGreen (Biotium Inc), and imaged with a ChemiDoc MP imaging system (BioRad).

### Cloning of RPA

The RPA ORFs (Rfa1, Rfa2 and Rfa3) were amplified from *S. cerevisiae* genomic DNA (SK1 strain), using the following primers Rfa1_Tag_Fwd (CTGTTCCAGGGGCCCGGATCC ATGAGCAGT-GTTCAACTTTCGAGGGGCGAT), Rfa1_rev (TC-CTCTAGTACTTCTCGACAAGCTTTTATTAAGC-TAACAAAGCCTTGGATAACTCATCGGCAAG), Rfa2_Tag_Fwd (CTGTTCCAGGGGCCCGGATC-CATGGCAACCTATCAACCATATAACGAATATTC), Rfa2_rev (TCCTCTAGTACTTCTCGACAAGCTTT-TATCATAGGGCAAAGAAGTTATTGTCATCAAAAG), Rfa3_Tag_Fwd (CTGTTCCAGGGGCCCGGATCCATG-GCCAGCGAAACACCAAGAGTTGACCCC), Rfa3_rev (TCCTCTAGTACTTCTCGACAAGCTTTTACTAG-TATATTTCTGGGTATTTCTTACATAG). Rfa1 was cloned into pCOLI_A_Strep, Rfa2 into pCOLI_K_His and Rfa3 into pCOLI_S_His. For the multicistronic assembly the Rfa1 RBS/ORF was amplified using the Alpha_Fwd and CasI_rev primers; Rfa2 with the CasII_fwd and CasII_rev primers and Rfa3 with CasIII_fwd and Omega_rev primers (Supplementary Table 2). The PCR amplified RBS/ORFs for each of the three RPA subunits were then assembled into linearized pCOLI_G2, with a 3-5 fold molar excess over the plasmid backbone, as previously described for pBIG assembly^5,6^. Gibson reactions were transformed directly into chemically competent XL1-blue *E. coli*, and selected on gentamycin LB agar plates. Minipreps of eight positive colonies were prepared, and subject to SwaI digest to release the individual RBS/ORF cassettes. Digests were then subject to agarose gel electrophoresis, and those clones that had bands of the appropriate molecular weight were sequence verified.

### Bacterial test expressions

Chemically competent BL21(DE3) E. coli were transformed with either a combination of pCOLI_A_Strep_Rfa1, pCOLI_K_His_Rfa2 and pCOLI_S_Rfa3 (co-transformation) OR pCOLI_G2_RPA (multicistronic assembly). 25 mL LB shake cultures of *E. coli* were grown in the presence of all appropriate antibiotics at 37°C. As the culture reached an OD600 of 0.6 IPTG was added to a final concentration of 500 µM, for a 3 hour induction. Cells were harvested, and resuspended in lysis buffer (50 mM Na-HEPES pH 7.5, 300 mM NaCl, 10% glycerol, 1 mM MgCl2, 2 mM BME, 1 mM AEBSF, 2.5 units/mL benzonase). Resuspended cells were then broken using sonication, and the lysate cleared by ultra-centrifugation. The cleared lysate was subject to affinity purification using Strep-Tactin XT resin (IBA), according to the manufacturer’s instructions. The resin was subject to several washes with ice-cold lysis buffer, before elution with lysis buffer supplemented with biotin. Fractions from the expression/purification were analysed using SDS-PAGE stained with InstantBlue (Sigma). Western blotting was carried out using standard laboratory protocols, using the anti-PentaHis (Qiagen) and anti-Strep II (Abcam ab76949) as primary antibodies and HRP conjugated anti-mouse or anti-rabbit secondary antibodies (Merck).

## Supporting information

Vector Sequence Files

## ACKNOWLEDGEMENTS

We thank the members of the Weir Lab for extensive testing of the system, and for comments on the manuscript. Work in the Weir Lab is funded by the Max Planck Society.

---

**Supplementary Table 1:**
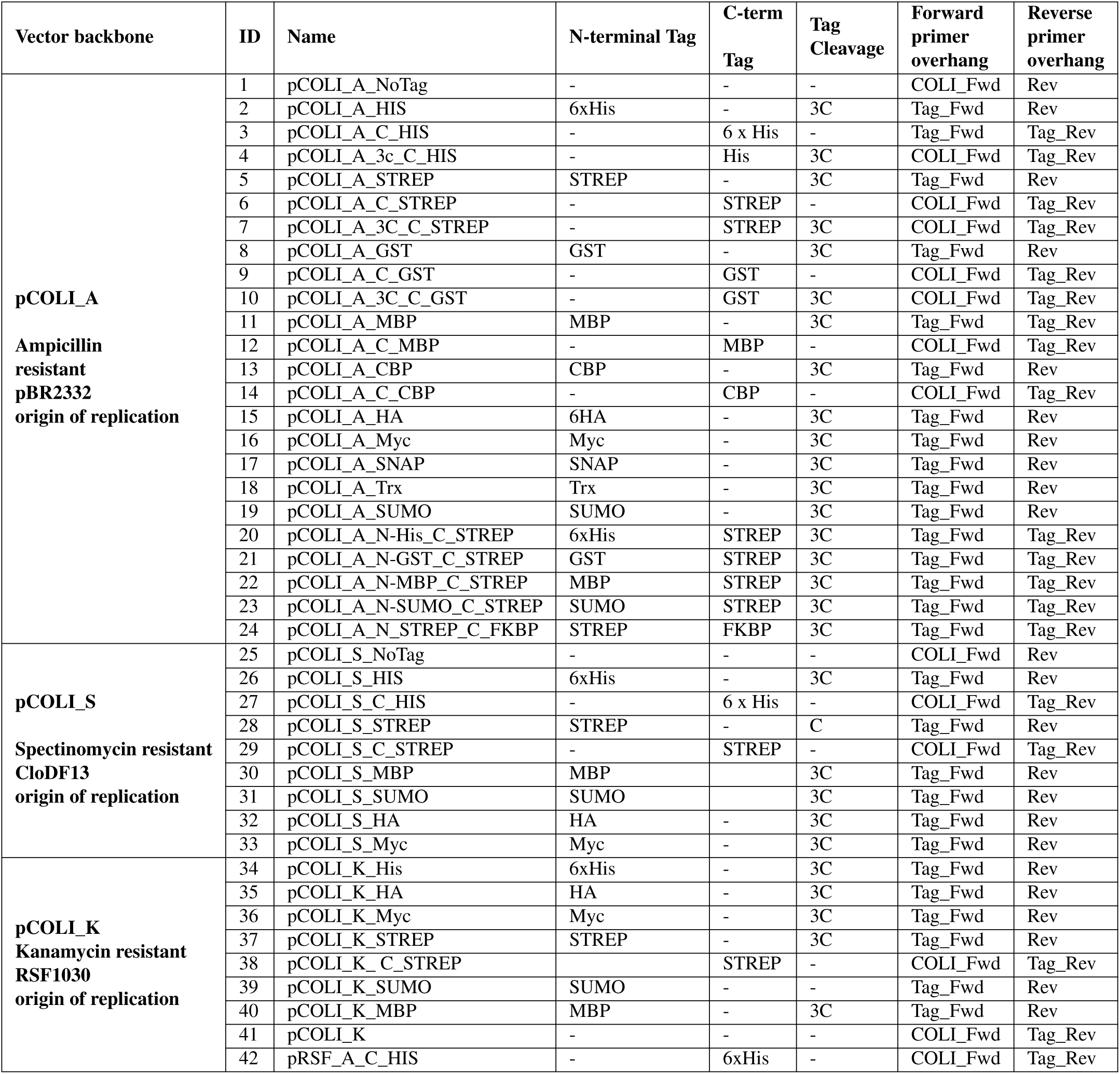
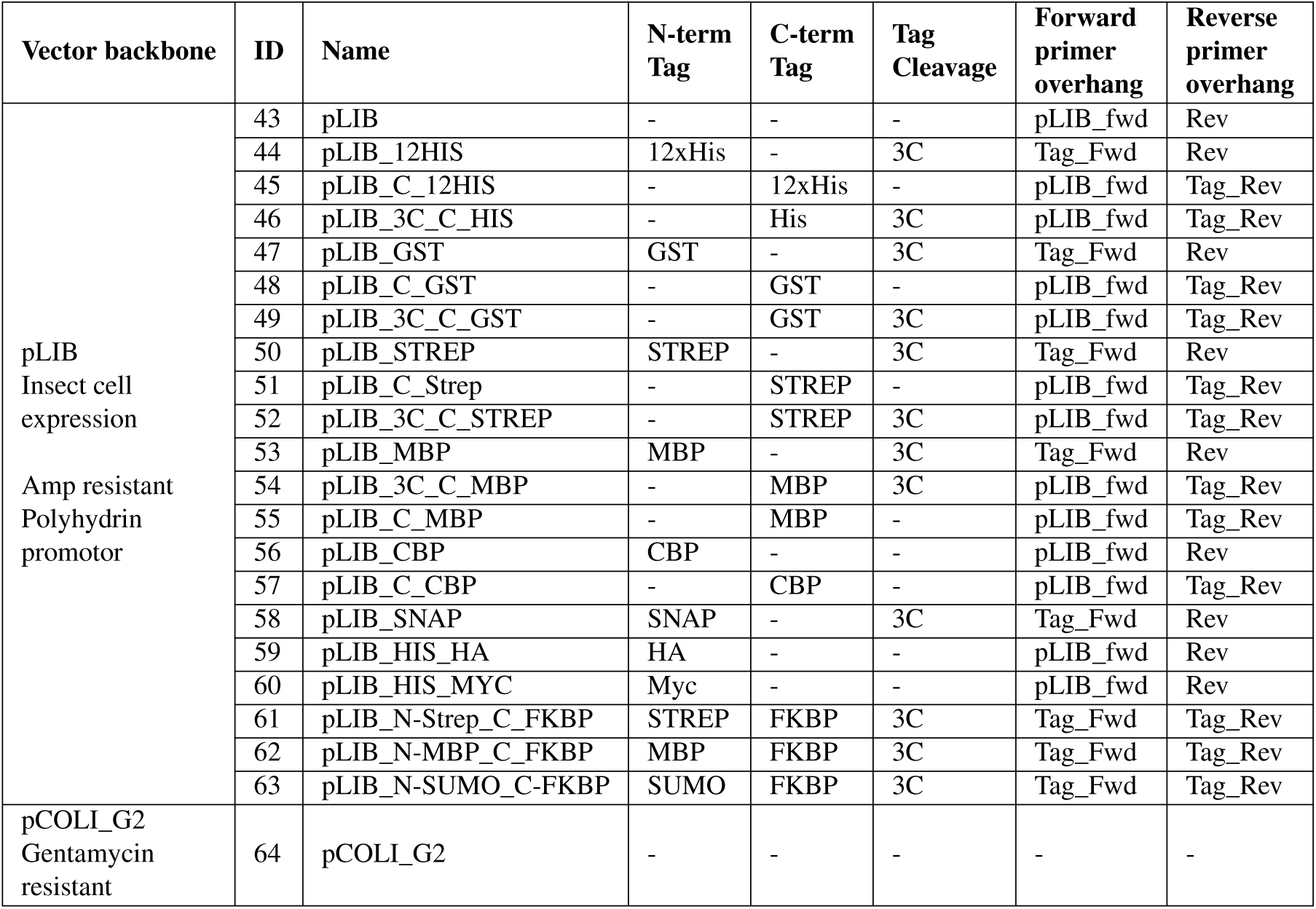
InteBac Vector Suite.

**Supplementary Table 2:**
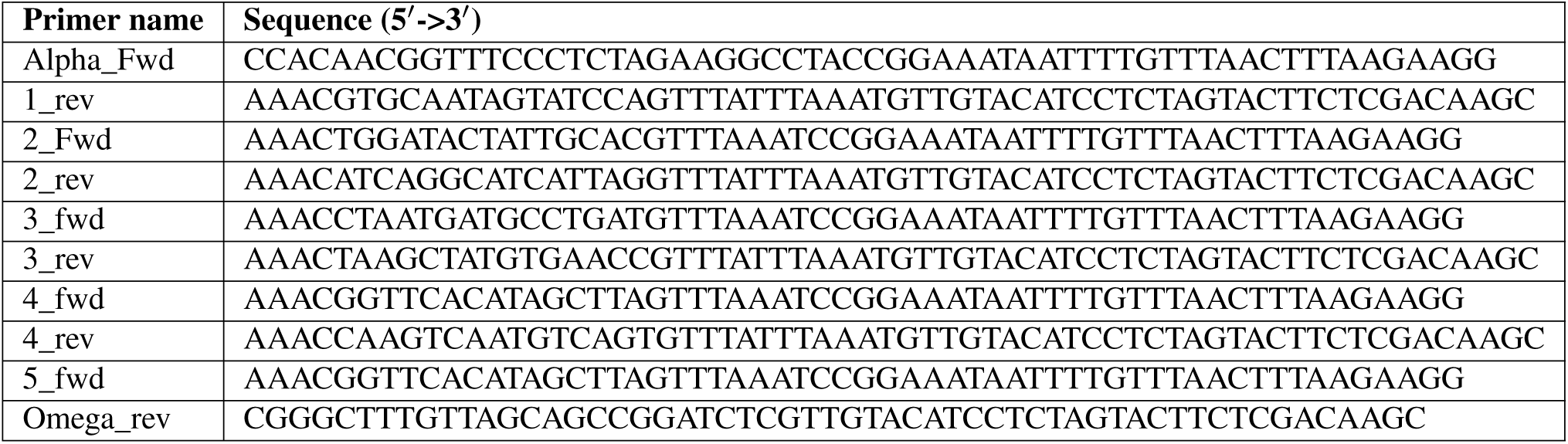
Primers for Multicistronic cloning into pCOLI_G2.

